# Experimental evolution of *S. cerevisiae* for caffeine tolerance alters multidrug resistance and TOR signaling pathways

**DOI:** 10.1101/2024.04.28.591555

**Authors:** Renee C. Geck, Naomi G. Moresi, Leah M. Anderson, yEvo Students, Rebecca Brewer, Timothy R. Renz, M. Bryce Taylor, Maitreya J. Dunham

## Abstract

Caffeine is a natural compound that inhibits the major cellular signaling regulator TOR, leading to widespread effects including growth inhibition. *S. cerevisiae* yeast can adapt to tolerate high concentrations of caffeine in coffee and cacao fermentations and in experimental systems. While many factors affecting caffeine tolerance and TOR signaling have been identified, further characterization of their interactions and regulation remain to be studied. We used experimental evolution of *S. cerevisiae* to study the genetic contributions to caffeine tolerance in yeast, through a collaboration between high school students evolving yeast populations coupled with further research exploration in university labs. We identified multiple evolved yeast populations with mutations in *PDR1* and *PDR5*, which contribute to multidrug resistance, and showed that gain-of-function mutations in multidrug resistance family transcription factors *PDR1, PDR3*, and *YRR1* differentially contribute to caffeine tolerance. We also identified loss-of-function mutations in TOR effectors *SIT4, SKY1*, and *TIP41*, and show that these mutations contribute to caffeine tolerance. These findings support the importance of both the multidrug resistance family and TOR signaling in caffeine tolerance, and can inform future exploration of networks affected by caffeine and other TOR inhibitors in model systems and industrial applications.

## Introduction

Caffeine is an alkaloid produced by plants including coffee and cacao. It is desirable for its effects as a stimulant, but also inhibits the master signaling regulator TOR (target of rapamycin) (Reinke *et al*. 2006). *S. cerevisiae* yeast are used to ferment coffee and cacao pulp (Ludlow *et al*. 2016), and there they are exposed to concentrations of caffeine ranging from approximately 20-80mM (3.5-16mg/g) (Pandey *et al*. 2000; Asfew and Dekebo 2019; Melo *et al*. 2021). Thus *S. cerevisiae* is capable of growing in high concentrations of caffeine, though laboratory and baking strains have much lower caffeine tolerance (Peter *et al*. 2018; Moresi *et al*. 2023).

Previous experimental evolution of haploid *S. cerevisiae* established that increased caffeine tolerance in a laboratory strain can be selected for in 48 passages; the one sequenced evolved clone from this study had mutations in *PDR1, PDR5*, and *RIM8*, indicating a role for the multidrug resistance (MDR) pathway in caffeine tolerance (Sürmeli *et al*. 2019). Caffeine can be exported from yeast cells by MDR transporters Pdr5 and Snq2 (Tsujimoto *et al*. 2015), which are regulated by transcription factors Pdr1, Pdr3, and Yrr1 (Cui *et al*. 1998). Pdr5 and Snq2 are highly homologous, but Snq2 has higher activity for caffeine efflux (Tsujimoto *et al*. 2015).

We also previously conducted experimental evolution of diploid *S. cerevisiae* for fewer than 20 passages, and found evolved clones had increased caffeine tolerance and mutations in many genes including *PDR1* (Moresi *et al*. 2023). However, we did not identify any genes with mutations in multiple evolved populations, which made differentiation of causative versus passenger mutations and prioritization of follow-up studies beyond *PDR1* challenging. Work remains to more fully characterize inputs and regulators of TOR in yeast, so studying mutations that increase resistance to TOR inhibitors such as caffeine could identify additional sites, domains, and relationships that are critical for TOR signaling. Many components of TOR signaling are conserved across eukaryotes (González and Hall 2017), so studying TOR regulation in yeast can also help us understand conserved processes and identify differences across species.

In order to identify more genes and processes that contribute to caffeine tolerance, we performed experimental evolution of haploid *S. cerevisiae* to increase our ability to see effects of loss-of-function and recessive mutations. As in our prior study on diploid yeast, we conducted experimental evolution as part of yEvo (“yeast Evolution”), an authentic research experience for high school students (Taylor *et al*. 2024). Previous yEvo studies on the effects of the antifungal drug clotrimazole identified new mutations, genes, and pathways involved in clotrimazole resistance (Taylor *et al*. 2022). Through yEvo we were able to generate 45 independently evolved populations and involve high school students in scientific research. By studying the effects of mutations in caffeine-evolved yeast, we identified crucial roles for specific members of the multidrug resistance family and downstream effectors of TOR signaling.

## Materials and Methods

### Yeast strains and media

All evolution experiments were performed with MATα S288C derivative BY4742. All strain genotypes are listed in Table S1, plasmids in Table S2, and primers and oligos in Table S3. Strains used in evolution experiments carried a 2µm plasmid with KanMX, which provides resistance to the general antibiotic G418, and a pigment production pathway that gives each strain a unique color (courtesy of the Boeke lab at New York University); see Taylor et al. 2022 for additional description. All experiments unless otherwise noted were performed in YPD (10g yeast extract, 20g peptone, and 20g dextrose per liter). Caffeine (Sigma-Aldrich) was added to YPD for a final concentration of 40mM. G418 disulfate (KSE Scientific, prepared as 200mg/mL stock in water) was added to the media (200 mg/L) for strains carrying pigment plasmids.

### Experimental evolution

Evolution experiments were carried out via batch transfer in YPD+G418 media. 1-2 times per week yeast were transferred using a sterile cotton swab from a saturated culture to a tube of 2-3mL fresh media containing caffeine. All experiments began in 10mM caffeine, and concentration was raised stepwise to 20, 30, and 40mM; the maximum concentration reached in each experiment is listed in Table S4. Evolved clones YMD4687-4712 were isolated from populations grown in 10-40mM caffeine for seven weeks in a 30°C incubator with no shaking, and YMD4821-4826 were grown in 10-40mM caffeine for eight weeks in a 30°C shaking incubator (BioRad, 70rpm). Evolved clones YMD4666-4686 were isolated from populations grown at room temperature in 10mM caffeine for five weeks; YMD4684-4686 were then grown at 20mM for an additional five weeks. At endpoint, glycerol stocks of each population were saved (1:1 yeast culture:50% glycerol), frozen at -20°C, and sent to the University of Washington for long-term storage at -72°C. Three clones were isolated from each population by streak purification and growth in caffeine measured as detailed below; one or two clones with the highest growth rate in the presence of caffeine were selected from each population for further experimentation. Clones isolated from the same population are noted in Table S5.

### Growth measurements

To assay growth in the presence of caffeine or clotrimazole, clones were grown in 200µL YPD medium in 96-well plates for 24 hours at 30°C. Cultures were resuspended and 2µL of each culture was transferred to 198µL YPD + indicated concentration of caffeine or clotrimazole. For evolved clones and ancestors, which contain pigment plasmids, 200mg/L G418 was also added. Growth of these cultures was monitored in a BioTek Synergy H1 microplate reader for 48 hours at 30°C with orbital shaking. The average growth rate of all strains with the same genotype was calculated by linear fit to logarithmic growth phase with R-squared cutoff of 0.85 and plotted in R; script is available on GitHub (github.com/reneegeck/DunhamLab/blob/main/platereader_growthplotter.R).

For growth with rapamycin, clones were grown overnight in 2mL YPD medium, with 200mg/L G418 for strains containing pigment plasmids, at 30°C in a roller drum. This larger culture format was used because when in 96-well cultures, even high concentrations of rapamycin (>1µM) did not fully inhibit growth of ancestor strains, in contrast to prior reports that 25nM leads to growth arrest (Barbet *et al*. 1996); growth in the larger cultures more closely matched expected effects. Experiment cultures were started at OD=0.05 in 2mL YPD -/+ 5nM rapamycin (LC Laboratories, 10mg/mL stock in ethanol), +200mg/L G418 if carrying pigment plasmid, and grown for 24 hours at 30°C in a roller drum. Optical density at 600nM was measured on a BioRad SmartSpec 3000.

To assay petite status, growth on YPD agar was observed, and clones were additionally transferred to YPG agar plates (glycerol as carbon source) and presence or absence of growth noted after 48 hours at 30°C.

### Whole-genome sequencing

Sample preparation, sequencing, determination of copy number variation, and identification of SNPs and indels was performed as previously described (Taylor *et al*. 2022). Briefly, DNA was purified using a phenol-chloroform and ethanol precipitation methodology. This DNA was tagmented using a modified Illumina Tagmentation protocol (Baym *et al*. 2015). Evolved clones were sequenced on an Illumina NextSeq 550 to an average coverage of 75X (range 10-178). Variants present in the ancestral strain YMD4612 (sequenced to 165X) were removed from consideration. Transposable element insertions were identified using McClintock v.2.0.0 (Nelson *et al*. 2017), filtered for non-ancestral insertions, and inspected if they were identified by at least three component methods per clone. All point mutations, copy number variations, and transposable element insertions were manually inspected in the Integrative Genomics Viewer (Robinson *et al*. 2011) to ensure they were correctly called and not present in the ancestor. Mutations, transposon insertions, and copy number alterations for each clone are listed in Table S5. Sequencing reads are deposited in the NCBI Sequence Read Archive (SRA) under BioProject PRJNA1101923.

### CRISPR-Cas9 construction of point mutations

The nearest NGG PAM (polyspacer adjacent motif) sequence to the mutation of interest was manually identified, and confirmed that it could be mutated by a synonymous mutation to stop Cas9 from cutting after a successful edit. Guide RNA (gRNA) sequence was selected as 20bp upstream of the PAM. Oligos (Table S3) were ordered from Integrated DNA Technologies to assemble the gRNA sequence into pNA0525 (Lauer *et al*. 2023) at the NotI cut site. Oligos were annealed and filled by two cycles of PCR, 5ng assembled into 90ng NotI-digested pNA0525 using Gibson Assembly Master Mix (New England Biosciences E2611), transformed into NEB5α *E. coli* (New England Biosciences C2987), and selected on LB+Ampicillin agar plates. Successful integration of gRNA was confirmed using Sanger sequencing (GENEWIZ from Azenta Life Sciences). Donor templates with 40bp of homology on each side of the desired edits were ordered and annealed as with gRNA. Strain YMD4042 was transformed with Cas9 expression plasmid pCTC019 (Zhao *et al*. 2023) and selected on c-leu to obtain strain YMD4909. For each mutation, an overnight culture of YMD4909 was transformed with 500ng pNA0525-gRNA and 500ng donor template using a lithium acetate protocol and selection on c-his-leu agar plates. Successful edits were confirmed using colony PCR and Sanger sequencing (Table S3).

### MDR transporter transcription reporters

Strains containing mutations in MDR transcription factors were transformed with a CEN plasmid containing *URA3* and *E. coli* gene lacZ under the control of *S. cerevisiae* promoters from *PDR5, SNQ2*, or *YOR1* (Katzmann *et al*. 1994; Kolaczkowska *et al*. 2008), or a control plasmid containing *URA3* but no lacZ gene (p416CYC1) (Mumberg *et al*. 1995). Resultant strains (YMD5073-86, YMD5138-49, Table S1) were grown overnight in c-ura, diluted to OD=0.1, and 5µL spotted onto c-ura agar plates containing 0.07M KPO_4_ pH7 and 40mg/L X-gal (American Biorganics, prepared as 2mg/mL stock in dimethylformamide). Plates were photographed after three days of growth at 30°C.

### Statistical analysis

All statistical analysis was performed in R. Code for statistical analysis and generating figures is available at https://github.com/reneegeck/DunhamLab/Geck2024_CaffeineYeast.

## Results

### Evolution of caffeine-tolerant yeast

Students and university lab members evolved 45 populations of *S. cerevisiae* in increasing concentrations of caffeine for five to ten weeks (approximately 10-20 transfers). Three clones were isolated from each population and tested for their ability to grow in the presence of caffeine relative to the ancestral strains; based on this, one or two clones per population were selected for further replicates of the growth assay (n=3 per clone) and downstream analysis for a total of 53 clones (Fig. 1, Table S4). The majority of clones (43/53, from 38/45 populations) had a significantly faster growth rate than the ancestor strains in caffeine (Fig. S1), indicating successful selection of caffeine-tolerant clones; some grew more slowly than the ancestor in the absence of caffeine, but not significantly slower (Fig. S1). Raising the concentration of caffeine in the evolution experiment selection was critical for selection of caffeine-tolerant clones, since 9/10 clones that did not grow better than the ancestor in caffeine were only grown in 10mM caffeine for the evolution process. Stepwise increasing the concentration also improved the ability to grow at higher concentrations of caffeine, but clones that only experienced 20-30mM caffeine during evolution still had significantly increased growth in 40mM caffeine (Fig. 1).

**Figure 1:**
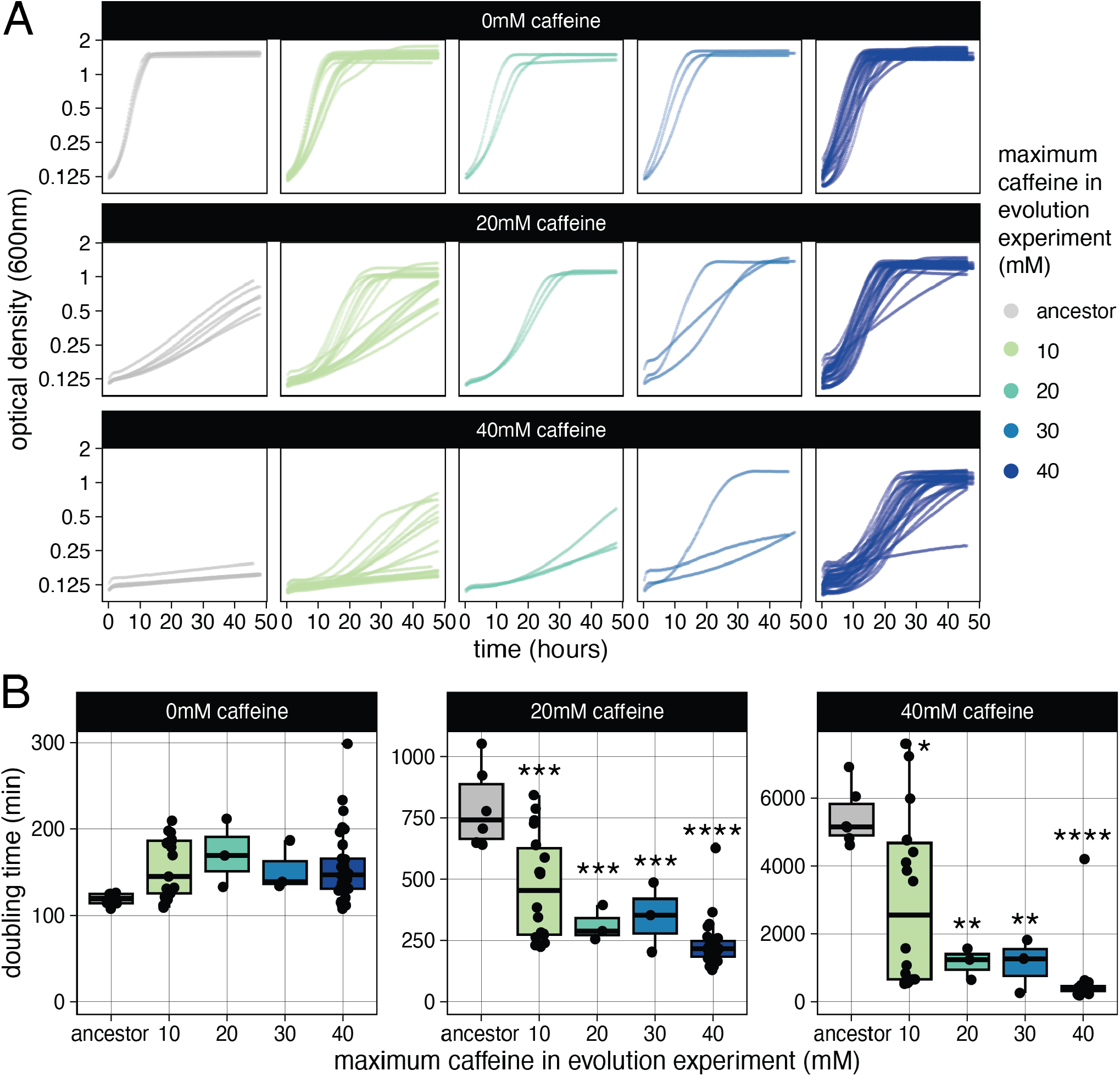
Evolution of caffeine tolerance. (A) Growth curves of ancestors and evolved clones. Each curve represents one clone and is the average of 3 biological replicates. (B) Doubling times for ancestors and evolved clones in A. Difference from ancestor by ANOVA with Tukey’s HSD, ^*^ p < 0.05, ^**^ p < 0.01, ^***^ p < 0.001, ^****^ p < 0.0001.

### Genetic characterization of caffeine-evolved yeast

To investigate the genetic changes contributing to increased caffeine tolerance, we sequenced all 53 evolved clones. Of the ten clones without increased caffeine tolerance, most either had no mutations or only one mutation in a gene not previously found connected to caffeine or TOR (Table S6); three that lost mtDNA but lacked nonsynonymous mutations did grow slightly faster in caffeine than ancestor strains (Fig. S2A), in line with previous reports that loss of mtDNA has been previously shown to increase growth in the presence of caffeine (Bard *et al*. 1980). These ten clones were excluded from further analysis to focus only on mutations likely to contribute to caffeine tolerance.

Every caffeine-tolerant clone had at least one nonsynonymous point mutation, 17/43 also lost mitochondrial DNA (mtDNA), and 7/43 also had a Ty insertion or focal copy number gain (Fig. 2A, Table S5). Only two evolved clones had copy number alterations (Fig. S2B). Clone YMD4666 did not have increased caffeine tolerance (Fig. S1); clone YMD4679 had a duplication of chrIII:152000-167000. Several genes fall within this region, but one, the nucleotide pyrophosphatase *NPP1*, has been previously connected to caffeine tolerance through deletion set screens where *npp1Δ* strains were more sensitive to caffeine (Dudley *et al*. 2005; Brown *et al*. 2006).

**Figure 2:**
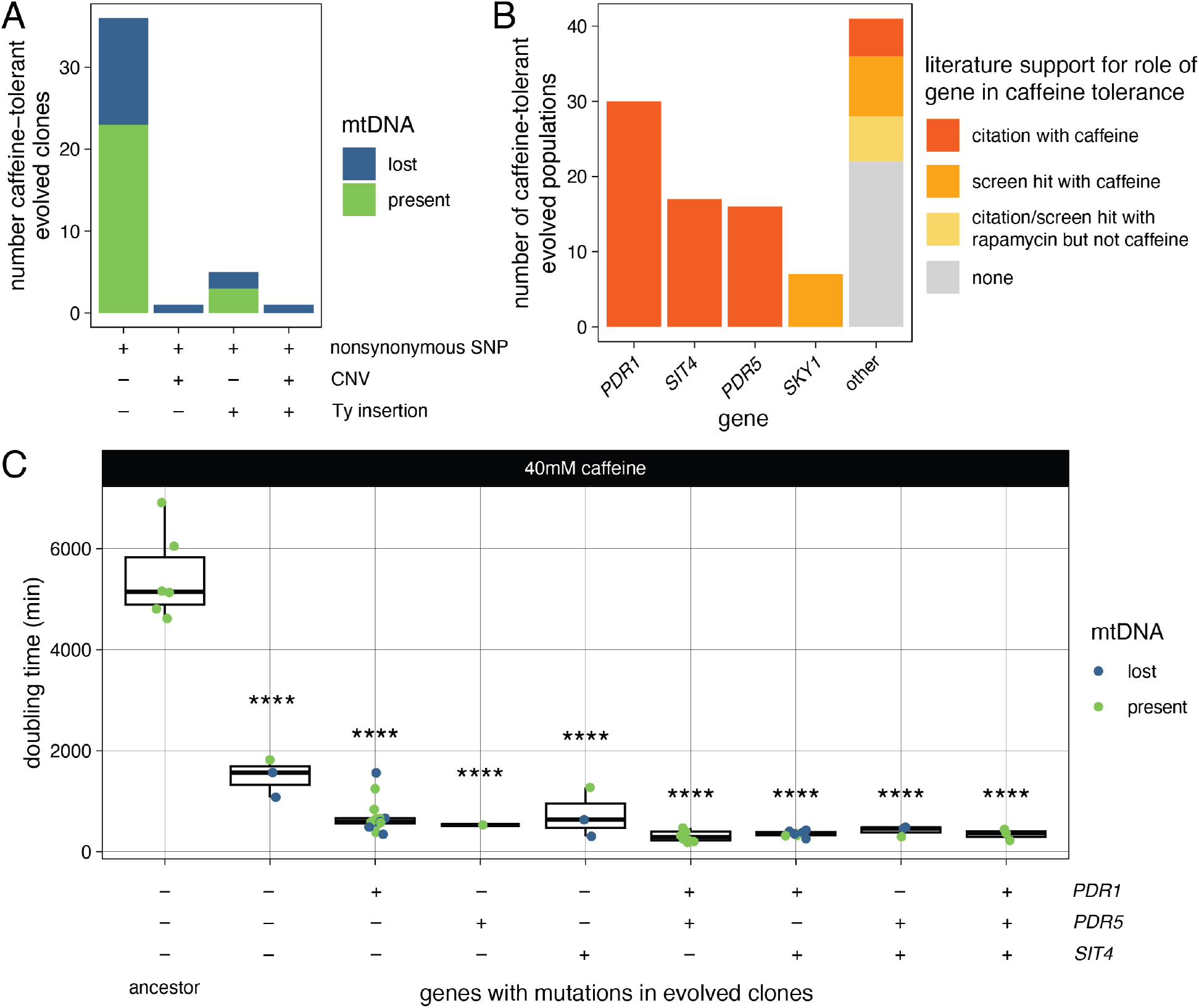
Common mutations in caffeine-tolerant clones. (A) Types of mutations in evolved clones with significantly increased caffeine tolerance. (B) Genes with mutations in caffeine-tolerant evolved lineages and their prior support for connection to caffeine tolerance or TOR signaling. (C) Doubling time of caffeine-tolerant clones grouped by mutations in multidrug resistance genes. Difference from ancestor by ANOVA with Tukey’s HSD, ^****^ p < 0.0001.

The nonsynonymous mutations were in 45 different genes (Table S6), 23 of which had been previously connected to caffeine or TOR signaling, including the only recurrently mutated genes *PDR1, PDR5, SIT4*, and *SKY1* (Fig. 2B, Table S7). We identified one mutation in *TOR1*, a target of caffeine, in the FKBP-rapamycin-binding (FRB) domain (*TOR1*^I1954M^). Another mutation at this same residue (*TOR1*^I1954V^) was previously shown to confer increased caffeine tolerance (Reinke *et al*. 2006).

The recurrently mutated genes *PDR1, PDR5*, and *SIT4* are involved in multidrug resistance (MDR) (Miranda *et al*. 2010; Buechel and Pinkett 2020), and all but three caffeine-tolerant evolved clones had a mutation in one or more of these genes (Fig. 2C). Mutations in *PDR1, PDR5*, and *SIT4* were found in populations evolved by slightly varied strategies in different classrooms (see Methods), emphasizing that the experimental evolution strategy is robust to variation. We did not have enough evolved clones in each group to determine if the effects of different combinations of *PDR1, PDR5*, and *SIT4* mutants had different effects on growth in caffeine, except that ones with mutations in both *PDR1* and *PDR5* grew faster in caffeine than ones with only *PDR1* mutations (Fig. S2C). The three clones without mutations in these genes (YMD4678, YMD4679, YMD4705) contained nonsynonymous mutations in *CAT8, SKY1, TIP41*, and *YRR1*. Mutations in *SKY1*, a downstream effector of TOR, were found in seven evolved populations (Fig. 2B). This included a nonsense mutation *SKY1*^Q621X^, indicating that *SKY1* loss of function is likely an important contributor to caffeine tolerance. This is supported by previous high-throughput findings that *sky1Δ* increases caffeine and rapamycin tolerance; Sky1 is also phosphorylated dependent on TOR (Brown *et al*. 2006; Huber *et al*. 2009).

### Cross-resistance conferred by mutations in MDR family genes

We hypothesized that many of the MDR mutations in our caffeine-evolved clones were not specific to caffeine but could confer resistance to other drugs. We previously evolved and characterized yeast with resistance to the antifungal drug clotrimazole, many of which also had gain-of-function mutations in *PDR1* or amplification of *PDR5* (Taylor *et al*. 2022). To determine if any mutations found in caffeine-tolerant clones confer cross-resistance to an unrelated drug, we grew clones in varying concentrations of clotrimazole. In general, growth in clotrimazole did not correlate strongly with growth in caffeine, though the clones that grow best in clotrimazole all have mutations in *PDR1* or *PDR5* (Fig. S3A-B, Table S8).

In our previous study of clotrimazole-evolved clones, we observed many mutations affecting *PDR1, PDR3*, and *PDR5*, but never *PDR1* and *PDR3* mutations in the same clone (Fig. S3C).

In contrast, a higher proportion of caffeine-evolved clones had point mutations in both *PDR1* and *PDR5*, whereas *PDR5* amplification was seen in clotrimazole-evolved clones alone or together with a *PDR1* or *PDR3* mutation. Additionally, while many caffeine-tolerant clones had *PDR1* mutations, none had a mutation in *PDR3*. Like the mutations in *PDR1* and *PDR3* from clotrimazole-evolved yeast, *PDR1* mutations in the caffeine-tolerant yeast were also gain-of-function since they also increased caffeine tolerance in a heterozygous diploid (Fig. S3D, Table S9). To investigate why we did not find any mutations in *PDR3* in our caffeine-evolved yeast, we grew clotrimazole-evolved yeast with gain-of-function mutations in *PDR1, PDR3*, or both in caffeine. When grown in clotrimazole, *PDR1*^F479I^ and *PDR3*^T949A^ confer equal resistance (Taylor *et al*. 2022). However, *PDR3* mutations were significantly less able to confer increased caffeine tolerance than *PDR1* mutations (Fig. 3A, S3E, Table S10). Therefore, most mutations in MDR genes conferred general drug resistance, but some had stronger effects on clotrimazole resistance than caffeine tolerance.

**Figure 3:**
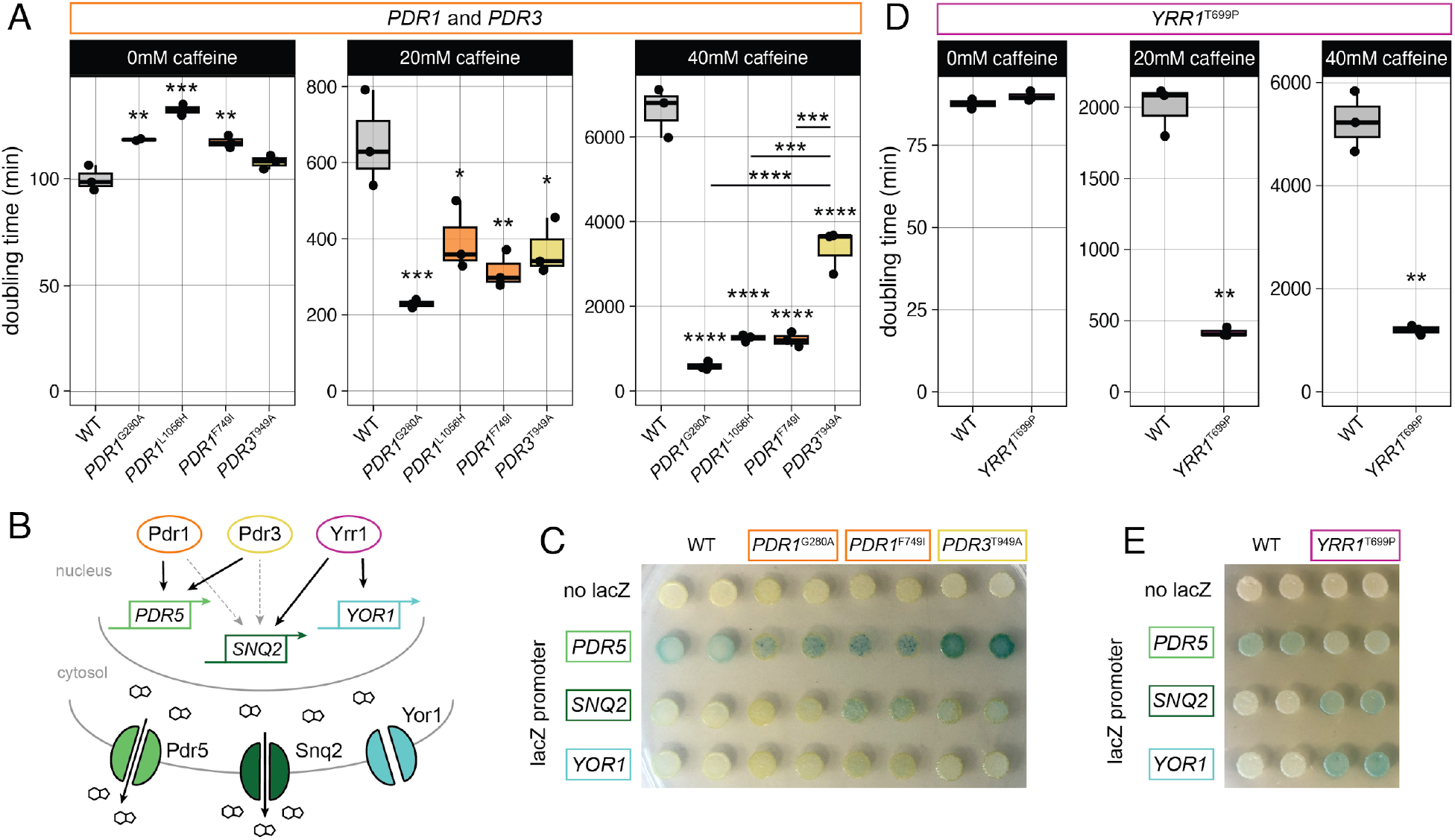
Gain-of-function mutations in MDR family genes contribute to cross-resistance to clotrimazole. (A) Doubling time of WT (YMD5096) compared to clones with *PDR1* and *PDR3* mutations from caffeine (YMD4681, YMD4684) or clotrimazole evolution experiments (YMD5097, YMD5098). (B) MDR transporters and their regulation by transcription factors based on Cui et al. 1998 and Kolaczkowska and Goffeau 1999, with effect on caffeine efflux from Tsujimoto et al. 2015. Dashed line arrows indicate basal activity; solid arrows indicate drug-induced activity. (C) Reporter assay of *PDR1* and *PDR3* mutant strains expressing lacZ under the control of indicated promoters, grown 72 hours on media containing X-gal. (D) Growth in caffeine of CRISPR engineered strains with synonymous *YRR1*^T696=^ mutation, with or without *YRR1*^T699P^ mutation. (E) Reporter assay of engineered *YRR1* strains expressing lacZ under the control of indicated promoters, grown 72 hours on media containing X-gal. Difference from wild type or synonymous by ANOVA with Tukey’s HSD, ^**^ p < 0.01, ^***^ p < 0.001, ^****^ p < 0.0001.

### Pdr1, Pdr3, and Yrr1 regulate caffeine tolerance

The two main efflux transporters for caffeine are *SNQ2* and *PDR5* (Fig. 3B) (Tsujimoto *et al*. 2015). To determine which targets were most affected by the gain-of-function mutations in *PDR1* and *PDR3*, we used transcriptional reporters for *PDR5, SNQ2*, and *YOR1*. Each reporter contains the *E. coli* lacZ gene under control of the promoter of *PDR5, SNQ2*, or *YOR1* (Kolaczkowska *et al*. 2008), so if the transcription is increased, yeast colonies will turn blue when grown on media containing X-gal. We observed that *PDR1*^G202A^, *PDR1*^F749I^, and *PDR3*^T949A^ led to an increase in transcription from the *PDR5* promoter (Fig. 3C, S3F), in line with our conclusion that these mutations increase function since *PDR5* is a target of Pdr1 and Pdr3 (Kolaczkowska and Goffeau 1999), but only a small increase in transcription from the *SNQ2* promoter. However, this does not explain the difference in caffeine tolerance provided by *PDR1* mutations over *PDR3* mutations (Fig. 3A), since *PDR3*^T949A^ led to more transcription from *PDR5* and *SNQ2* promoters than *PDR1*^G202A^, contrary to *PDR1*^G202A^ conferring faster growth in caffeine.

We also wanted to investigate if any mutations in caffeine-tolerant yeast had a greater effect on *SNQ2* expression. While Pdr1 and Pdr3 are important for basal expression of *SNQ2*, increased activation in response to drugs is mainly mediated through another MDR transcription factor, Yrr1 (Cui *et al*. 1998) (Fig. 3B). *YRR1* was previously found to have increased expression in a caffeine-tolerant evolved yeast strain (Sürmeli *et al*. 2019), and we found a mutation in *YRR1* in one of our caffeine-tolerant clones (Table S6). We hypothesized that our observed *YRR1*^T699P^ mutation leads to increased Yrr1 activity and thus contributes to MDR. We constructed the *YRR1*^T699P^ mutation using CRISPR-Cas9, and observed that it was sufficient to increase growth in caffeine in a haploid (Fig. 3D, S3G, Table S11) and in a heterozygous diploid (Fig. S3H, Table S11). Transcription from the *PDR5* promoter was basally active, and the mutation did not increase it, but *YRR1*^T699P^ did increase transcription from *SNQ2* and *YOR1* promoters (Fig. 3E, S3F), supporting that this mutation increases Yrr1 activity towards its targets. This is a similar transcriptional effect as seen by other gain-of-function *YRR1* mutations (Kodo *et al*. 2013). Thus Pdr1, Pdr3, and Yrr1 activity are all important for caffeine tolerance, likely by their effects on caffeine efflux transporters Pdr5 and Snq2.

### Many loss-of-function mutations contribute to caffeine and rapamycin tolerance

To investigate mutations that are more specific to caffeine effects on TOR signaling, we also looked at cross-resistance to rapamycin. Caffeine and rapamycin both bind the FRB domain of Tor1 (Reinke *et al*. 2006), so mutations that confer caffeine tolerance by affecting TOR signaling should also confer rapamycin resistance. We confirmed that evolved clone YMD4711, which has a mutation in the FRB domain of Tor1, has increased growth in caffeine (Fig. S1), and its growth is not inhibited by rapamycin (Table S12). Across all strains, growth in caffeine weakly correlates with growth in rapamycin, though the majority of evolved clones are highly resistant to rapamycin and double rapidly in caffeine (Fig. S4A, Table S12).

Two of the recurrently mutated genes, *SIT4* and *SKY1* (Fig. 2B), have been previously shown to affect rapamycin tolerance (López-Mirabal *et al*. 2008; Huber *et al*. 2009). We identified likely loss-of-function mutations in both genes in caffeine-tolerant clones that did not have mutations in *PDR1* or *PDR5*, suggesting the *SIT4* and *SKY1* loss-of-function may be conferring caffeine tolerance. To investigate more genes that may be implicated in caffeine and rapamycin resistance, we focused on all genes in caffeine-tolerant strains with likely loss-of-function mutations (*EBS1*^I18indel^, *ESL1*^W658X^, *SIT4*^N269indel^, *SKY1*^Q621X^, *SVF1*^F284indel^, *TIP41*^S162X^, *TOM20*^S36indel^, *USA1*^G37indel^). Of these nine genes, *ESL1* is the only essential gene, but hypomorphic alleles have been constructed (Breslow *et al*. 2008), so *ESL1*^W658X^ is likely hypomorphic. Deletions of the other genes were available in the yeast deletion collection (Tong *et al*. 2001; Pan *et al*. 2004), so we measured their growth in caffeine and rapamycin. Deletions of *EBS1, ESL1, SIT1, SKY1, SVF1, TIP41*, and *USA1* led to increased growth in caffeine, indicating that the loss-of-function mutations we observed in these genes are not passengers from the evolution experiment (Fig. 4A, S4B, Table S13). Only *sit4Δ, svf1Δ* and *tip41Δ* also increased growth in rapamycin (Fig. 4B, Table S14), indicating that they contribute to caffeine tolerance through TOR signaling, and that the role of Sit4 in caffeine tolerance may be through both the MDR pathway and TOR signaling (López-Mirabal *et al*. 2008; Miranda *et al*. 2010). However, in a previous screen of the systematic deletion set *svf1Δ* increased sensitivity to rapamycin (Xie *et al*. 2005), so its effects may be dose- or assay-dependent. There is also literature support that *ebs1Δ* and *sky1Δ* confer some resistance to rapamycin (Ford *et al*. 2006; Huber *et al*. 2009), suggesting our assay may not capture smaller effects.

**Figure 4:**
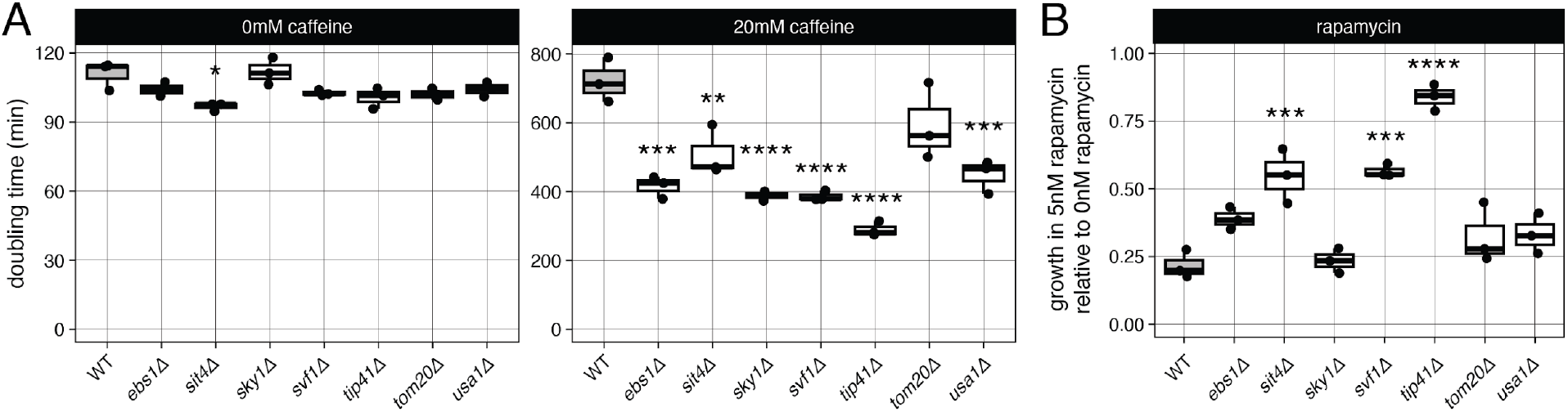
Effect of deletions on caffeine and rapamycin tolerance. (A) Doubling time and (B) growth after 24 hours in rapamycin for strains with indicated deletions. Difference from wild type (WT) by ANOVA with Tukey’s HSD, ^*^ p < 0.05, ^**^ p < 0.01, ^***^ p < 0.001, ^****^ p < 0.0001.

### TOR effectors Sky1, Sit4, and Tip41 contribute to caffeine tolerance

*SIT4* and *TIP41* are involved in PP2A-like signaling downstream of TOR, as is *RRD1*, in which we found a *RRD1*^D143H^ mutation (Fig. 5A). Loss of *SIT4* and *RRD1* have been previously shown to increase caffeine tolerance in high-throughput screens (Kapitzky *et al*. 2010; Hood-DeGrenier 2011), but *TIP41* has not. To confirm the contribution of *TIP41*^S162X^ we constructed the mutation using CRISPR-Cas9, and observed that *TIP41*^S162X^ increased growth in caffeine and rapamycin (Fig. 5B-C, S5A, Tables S11, S14). The *TIP41*^S162X^ mutation had no effect on caffeine tolerance in a heterozygous diploid, supporting that it is loss of function and recessive (Fig. S5B-C). We also constructed a point mutation in *SIT4, SIT4*^T95R^, which increased growth in the presence of caffeine and rapamycin (Fig. 5B,D, S5A, Tables S11, S14), indicating that inhibition of PP2A-like signaling is important for caffeine tolerance. The *SIT4*^T95R^ mutation is likely loss of function since it had no effect on caffeine tolerance in a heterozygous diploid (Fig. S5B-C).

**Figure 5:**
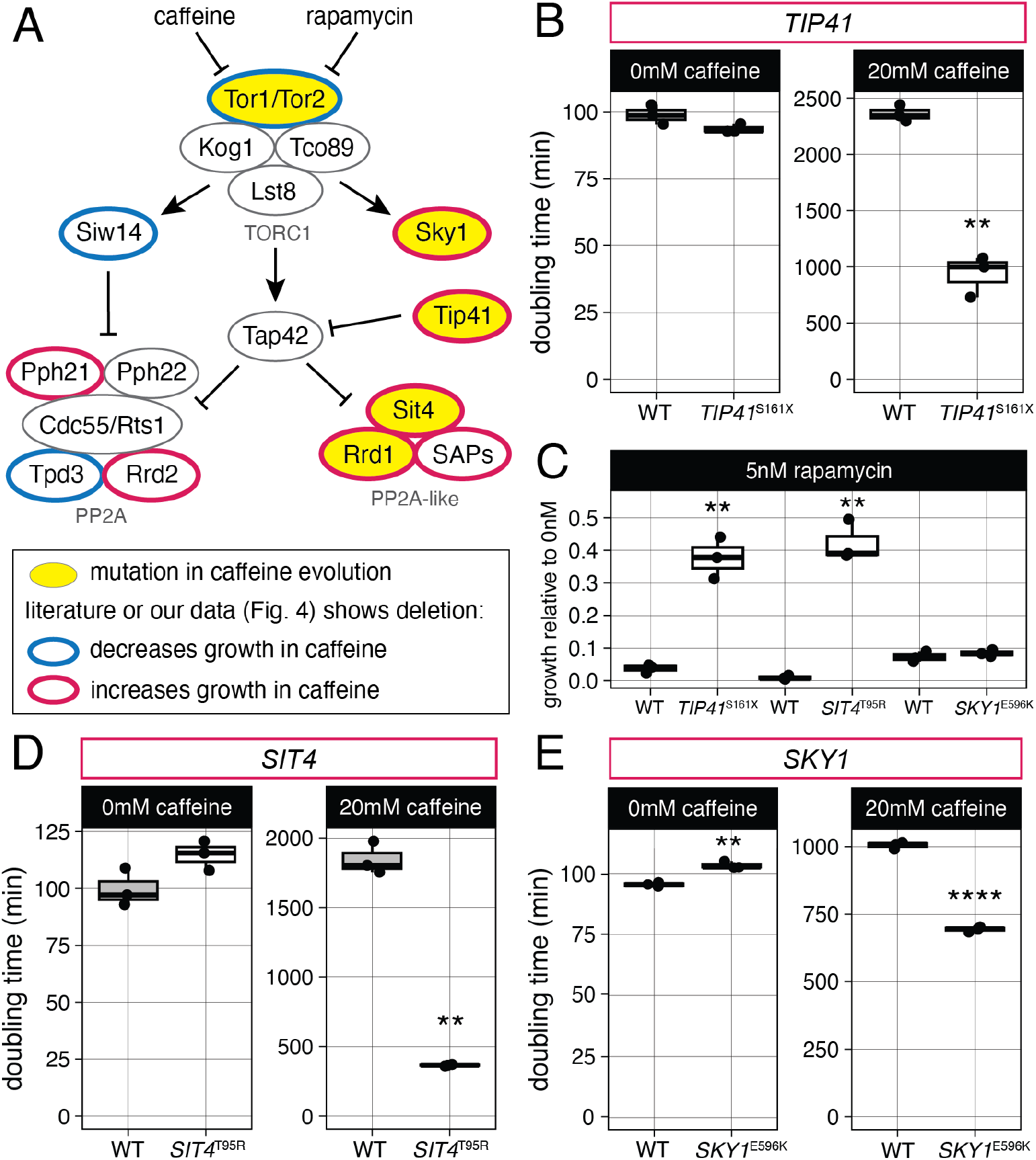
Contributions of TOR effectors to caffeine tolerance. (A) TOR signaling pathway (Huber *et al*. 2009; Numamoto *et al*. 2015; Ferrari *et al*. 2017; Ariño *et al*. 2019) annotated with mutations found in caffeine-tolerant clones in this study, and prior literature support for effects of deletions on caffeine tolerance (Rempola *et al*. 2000; Brown *et al*. 2006; Reinke *et al*. 2006; Banuelos *et al*. 2010; Kapitzky *et al*. 2010; Hood-DeGrenier 2011). (B) Doubling time for strains engineered to have synonymous *TIP41*^Q160=^, with or without *TIP41*^S161X^ mutation. (C) Growth after 24 hours in 5nM rapamycin for engineered strains. (D) Doubling time for strains engineered to have *SIT4* mutation; both have synonymous *SIT4*^L91=^ mutation. (E) Doubling time for strains with *SKY1* mutation; both have synonymous *SKY1*^T599=^ mutation. Difference from WT by t-test, ^**^ p < 0.01, ^****^ p < 0.0001.

Sky1 also functions downstream of TOR, but unlike Tip41 and Sit4 it directly associates with Tor1 instead of with Tap42 (Fig. 5A) (Huber *et al*. 2009; Breitkreutz *et al*. 2010). We constructed a *SKY1*^E596K^ point mutation using CRISPR-Cas9 and observed that it increased growth in caffeine in a haploid (Fig. 5G, S5A, Table S11) but not heterozygous diploid (Fig. S5B-C), indicating that the mutation likely decreases Sky1 function to increase caffeine tolerance. As seen with *sky1Δ, SKY1*^E596K^ did not significantly increase growth in 5nM rapamycin (Fig. 5B, Table S14).

## Discussion

We successfully evolved laboratory yeast to have increased caffeine tolerance, and identified contributing mutations in genes related to multidrug resistance and TOR signaling. In every caffeine-tolerant evolved clone, we identified at least one likely causative mutation: either in MDR pathway genes *PDR1, PDR5*, and *YRR1*, or TOR pathway genes *TOR1, SKY1, TIP41*, and *SIT4*. This is in agreement with a previous experimental evolution study in caffeine, where one clone was sequenced and found to have mutations in *PDR1, PDR5*, and *RIM1* (Sürmeli *et al*. 2019). In many genes, we identified multiple mutations in caffeine-tolerant clones, some affecting the same or nearby sites. Future work focusing on the effects of groups of mutations could elucidate important functional and structural properties of these proteins.

The majority of our caffeine-evolved yeast clones had a mutation in a MDR pathway gene, and characterizing these mutations provides insight into functional interactions and important regions of MDR proteins. We observed many mutations in *PDR1*, but none in *PDR3*. This could be because the main caffeine efflux transporter genes *PDR5* and *SNQ2* are more strongly regulated by Pdr1 than Pdr3, since *pdr1Δ* leads to a much greater decrease in *PDR5* and *SNQ2* transcripts than *pdr3Δ* (Mahé *et al*. 1996), though we did not observe strong activation of *SNQ2* transcription and saw similar activation of *PDR5* transcription by Pdr1 and Pdr3 gain of function mutants. More quantitative measurement of the effects of these mutations, and broader study of their effects on transcription of other genes beyond *PDR5* and *SNQ2*, are needed to identify possible reasons for the difference in effect strength of Pdr1 and Pdr3 on caffeine tolerance. We also added to the known gain-of-function mutations in *YRR1* that increase *SNQ2* expression and caffeine tolerance (Kodo *et al*. 2013; Rong-Mullins *et al*. 2018). Several of these mutations are near the activation domain of Yrr1 (Gallagher *et al*. 2014), including the *YRR1*^T699P^ mutant we identified, suggesting that mutations in this region are important for conferring resistance to various drugs. By characterizing mutations in MDR pathway genes, we are able to identify more gain-of-function mutants, which increase our understanding of functional domains in these proteins.

The high percentage of evolved clones with a mutation in a MDR gene somewhat hampered our ability to study TOR signaling using caffeine, since most clones had a mutation that could increase caffeine efflux, so the selection pressure for rewiring of intracellular signaling was not as strong. In the future, evolving a strain lacking key MDR components could lead to selection for different types of mutations, as done previously for evolution of rapamycin-resistant *pdr1Δ* yeast (Wride *et al*. 2014).

While we did not identify new relationships between TOR signaling effectors, we did show that loss of function of Tip41 increased caffeine tolerance. We also observed many mutations in components of the PP2A-like complex, but none in PP2A complex members. Few PP2A-like specific downstream processes have been annotated, as many are presumed to be shared with PP2A (Jiang 2006), but specific alpha-arrestins and factors involved in cell cycle progression and budding have been proposed to be downstream of PP2A-like signals (Fernandez-Sarabia *et al*. 1992; Watanabe *et al*. 2009; Talaia *et al*. 2017; Bowman *et al*. 2022). Different regulatory strength of PP2A and PP2A-like could also lead to increased tolerance or benefit of mutations in PP2A-like factors that spare processes proposed to be only downstream of PP2A, such as autophagy (Yorimitsu *et al*. 2009). Studying the differential regulation of these downstream processes could increase our understanding of how yeast responds to caffeine and other drugs and stressors that affect TOR signaling. Yeast that are used for coffee and cacao fermentation are exposed to high concentrations of caffeine, and studying natural variation in their TOR pathway components compared to other domesticated yeast could address how critical different factors are for growth in caffeine. Many domesticated yeast have been isolated and sequenced (Peter *et al*. 2018), but few from coffee and cacao have had their whole genomes sequenced which would be necessary for this further analysis (Ludlow *et al*. 2016).

A strength of our study was in the number of independent evolution experiments, and ability to compare it to other experiments carried out in the same yEvo system but under a different selective pressure. Firstly, we showed that caffeine tolerance in haploid yeast can be selected for in far fewer than 48 passages, as was previously done (Sürmeli *et al*. 2019), since some of our caffeine-tolerant clones were derived from populations that were passaged only 10 times. Additionally, by comparing our selection for caffeine tolerance to prior yEvo selections for clotrimazole resistance, we identified notable similarities and differences in the mutations that were selected for in each regime. Clotrimazole-evolved yeast exhibited copy number changes, including whole chromosome gains, and mutations in *PDR3* (Taylor *et al*. 2022), while copy number changes were rare (2/45 populations) and *PDR3* mutations absent in caffeine-evolved yeast. Most mutations that affected efflux transporter *PDR5* in clotrimazole-evolved clones were copy number gains (11/13), whereas caffeine-evolved clones had many *PDR5* point mutations. Since both of these experiments started with the same strains and used the same experimental evolution protocol of serial transfers, we can conclude that the differences are due to the differential ability of these mutations to confer caffeine tolerance compared to clotrimazole resistance. As we carry out more experiments through yEvo, we are compiling a rich dataset to investigate these differences and similarities between the effects of different selective pressures.

## Data Availability Statement

All sequencing data are available at the NCBI Sequence Read Archive (SRA) under BioProject PRJNA1101923. Strains and plasmids are available upon request. Code for statistical analysis and generating figures is available at https://github.com/reneegeck/DunhamLab/Geck2024_CaffeineYeast.

## Acknowledgements

This work would not be possible without the following yEvo students and members of the Dunham lab who conducted selection experiments: Isabel Addepalli, Deepti Aggarwal, Prisha Agnihotri, Ahlaam A. Ali, Clara J. Amorosi, Abhinav Anand, Ashna Atukuri, Thang Awi, Insiya Basrai, Hitha Bathala, Sarang Bhide, Benjamin B. Cantor, Jocelyn Cervantes, Tridib Chakraborty, James Champlin, Ameen Chbihi, Felicia Chen, Hayley Chenfang, Reagan Choi, Sebastian Chokka, Julian D’Souza, Vivek Dandu, Arkesh Das, Margrette Dawoud, Victoria Dong, Riya Dutta, Graeme Edoff, Cecelia Fan, Rena Foo, Liam T. Glanville, Cristian Golat, Suhavi Grewal, Faye Guan, Aarya Gurav, Aranav Gupta, Krish Gupta, Siya Gupta, Osman Hameed, Rhea Hede-Sakhardande, Nushaba Hossain, Youssef Ibrahim, Jemi Isaac, Udayvir Jalf, Medha Jasti, Amar Jazvin, Okichy Johnny Jr., Vidhi Kamat, Venya Kandula, Lekhana Katuri, Keabe E. Kebede, Om Khuperkar, Emily Kim, Rishi Konduru, Salimah Kyaw, Daniel Lee, Tian Syun Lin, Karen Luo, Jwan Magsoosi, Mlahat Mahmood, Ronald Brent F. Marzan, Noyonima Masud, Jessica Mathew, Ava Miciuda, Trevor Morey, Anagha Nair, Naveen Natarajan, Aahil Abdul Nazeer, Usoatua Levei P. Noa, Shashank Pagadala, Hamin Paik, John Palomino, Kush Parikh, Naisha Phadke, Michelle V. Phan, Britta Pingree, Neal Podhuturi, Arya Prasad, Sonia Puri, Sanjini Rajkumar, Ananya Ramanan, Elliot M. Russell, Zachary L. Saad, Magdalena Sabalsa Gaytan, Francis L. Salazar, Anjali Sanil, Neespruha Shah, Mustafa Sharba, Prihensha Sharma, Sophia Showman, Soyeon Showman, Heejin Shyn, Aryan Singh, Saakshi Sovani, Shreya Srugaram, Rachel Stroia, Sanjana Sunilkumar, Nihil Suthy, Asma Syed, Ruthesh Thavamani, Nitya Upadhye, Rebecca Varghese, Annie Wang, Cynthia Wang, Roger Wang, Miya A. Watson, Theresa Wei, Myra L. Woody, Nancy Yao, Tyler Yee, Chiann-Ling Cindy Yeh, Jungbin Yoon, Jiaying Zhou, Tianhui Zhu. We also thank Noah Fredstrom for assistance with sequencing; Sandra Pennington, Scarlett Counihan, Owen Burris, and Marisol Jimenez Garcia for yeast media and reagent support; Dennis Godin for assistance running McClintock; and members of the Dunham lab for feedback. Leo Pallanck’s lab (University of Washington) generously provided rapamycin, and Scott Moye-Rowley’s lab (University of Iowa) shared their lacZ reporter plasmids.

## Supplemental Material

**Supplemental Figure 1:**
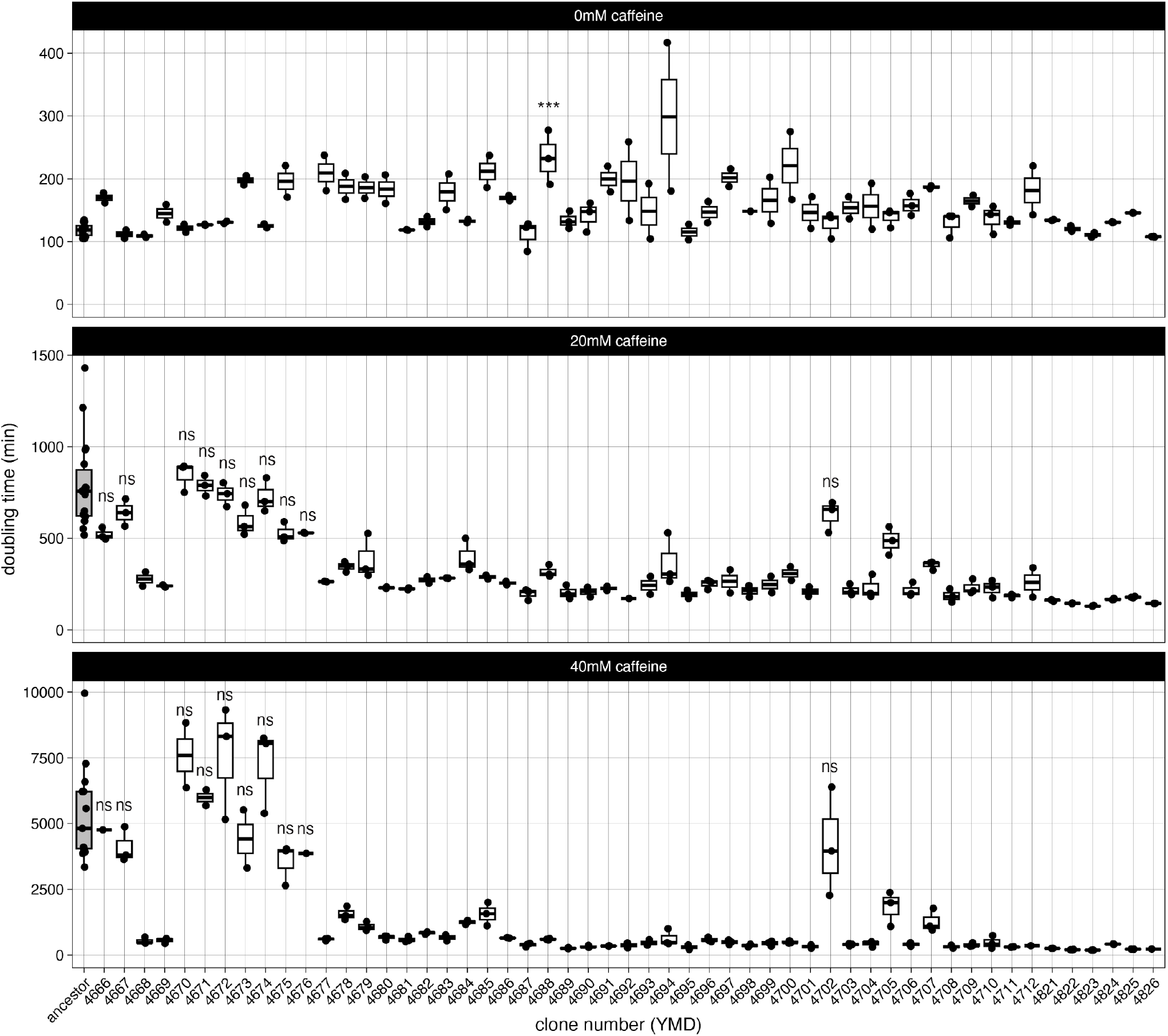
Doubling times for ancestors and caffeine-evolved clones. For 0mM caffeine, all differences from ancestors are *p* ≥ 0.05 by ANOVA with Tukey’s HSD, except YMD4688, *p* = 0.0004. For 20mM and 40mM caffeine, non-significant difference from ancestor denoted by “ns;” all other clones have *p* < 0.0001 except YMD4692 at 20mM (p = 0.0003), YMD4705 at 20mM (p = 0.018), YMD4712 at 40mM (p = 0.0002), and YMD4726 at 40mM (p = 0.0001).

**Supplemental Figure 2:**
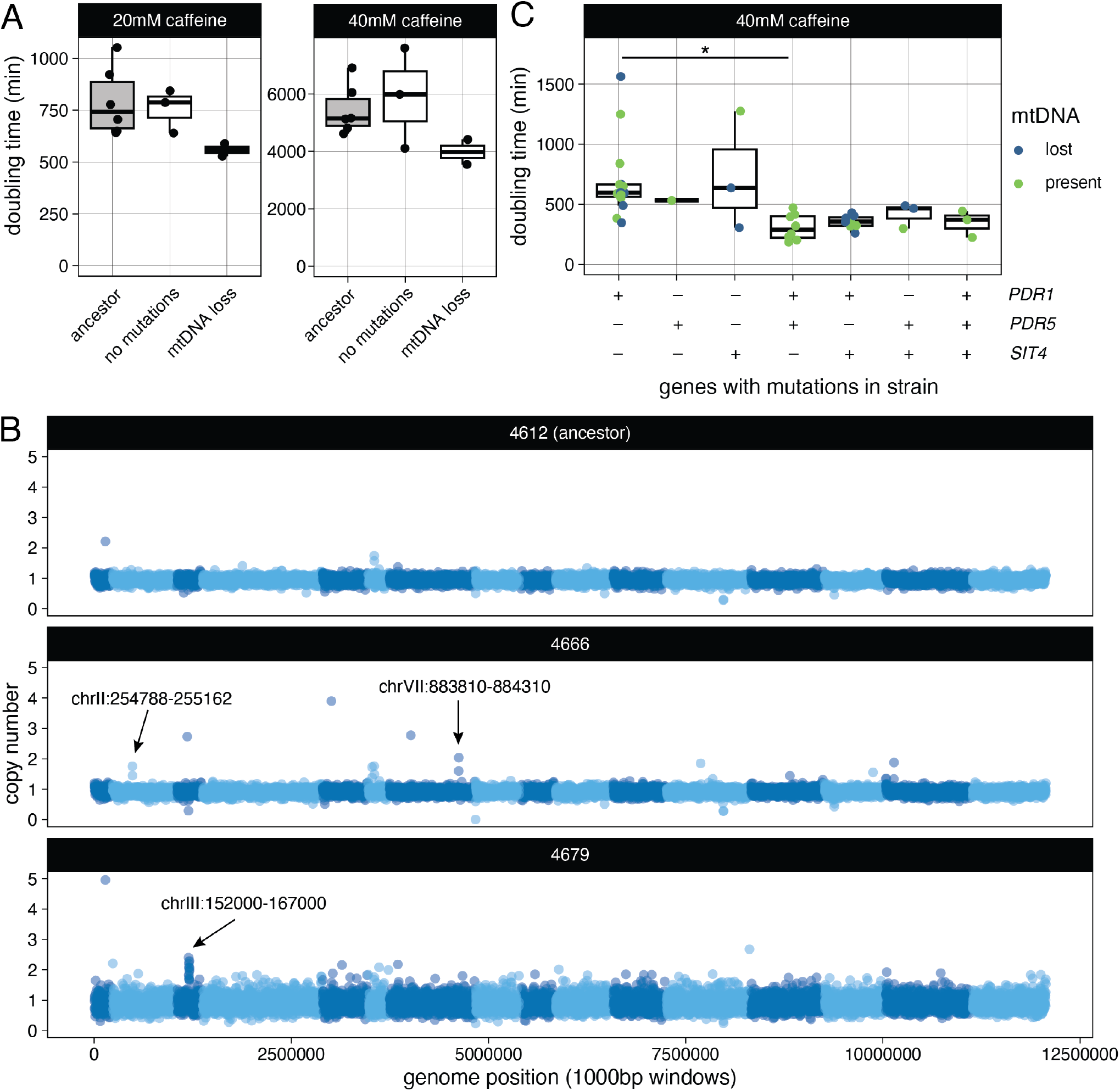
(A) Doubling times in 20mM and 40mM caffeine for ancestors and caffeine-evolved clones without nonsynonymous mutations. Differences are not statistically significant from ancestor by ANOVA with Tukey’s HSD. (B) Copy number inferred from read coverage of 1000bp windows. Arrows indicate alterations confirmed using IGV. (C) Doubling time of clones grouped by mutations in multidrug resistance genes; data from Fig. 2C without ancestor, none, and other groups. Difference between groups by ANOVA with Tukey’s HSD, ^**^ p < 0.01.

**Supplemental Figure 3:**
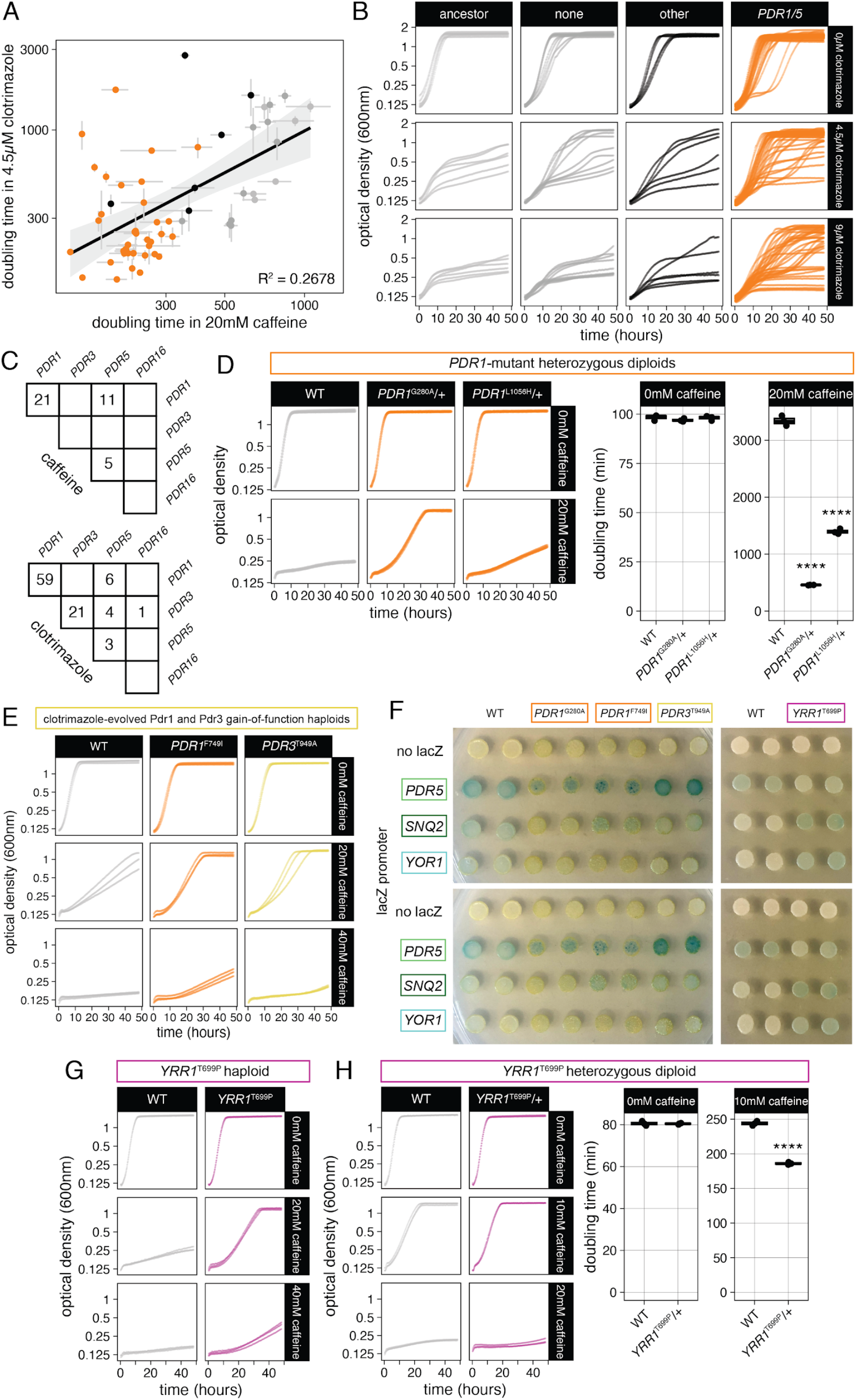
(A) Correlation between doubling time of caffeine-evolved clones and ancestors in caffeine and clotrimazole. (B) Growth curves of ancestors and caffeine-evolved clones in the presence of clotrimazole to test for cross-resistance. Each curve represents one clone and is the average of 3 biological replicates. None indicates evolved clones with no mutations detected. (C) Comparison of number of evolved clones with mutations, including *PDR5* amplification, in different PDR family genes from caffeine and clotrimazole experimental evolution. (D) Growth of diploid strains with heterozygous *PDR1* mutations. (E) Growth in caffeine of clones from clotrimazole evolutions with gain-of-function mutations in *PDR1* and *PDR3*. (F) Additional replicates of lacZ reporter assay. (G) Growth in caffeine of haploid or (H) heterozygous diploid CRISPR engineered strains with synonymous *YRR1*^T696=^ mutation, with or without *YRR1*^T699P^ mutation.

**Supplemental Figure 4:**
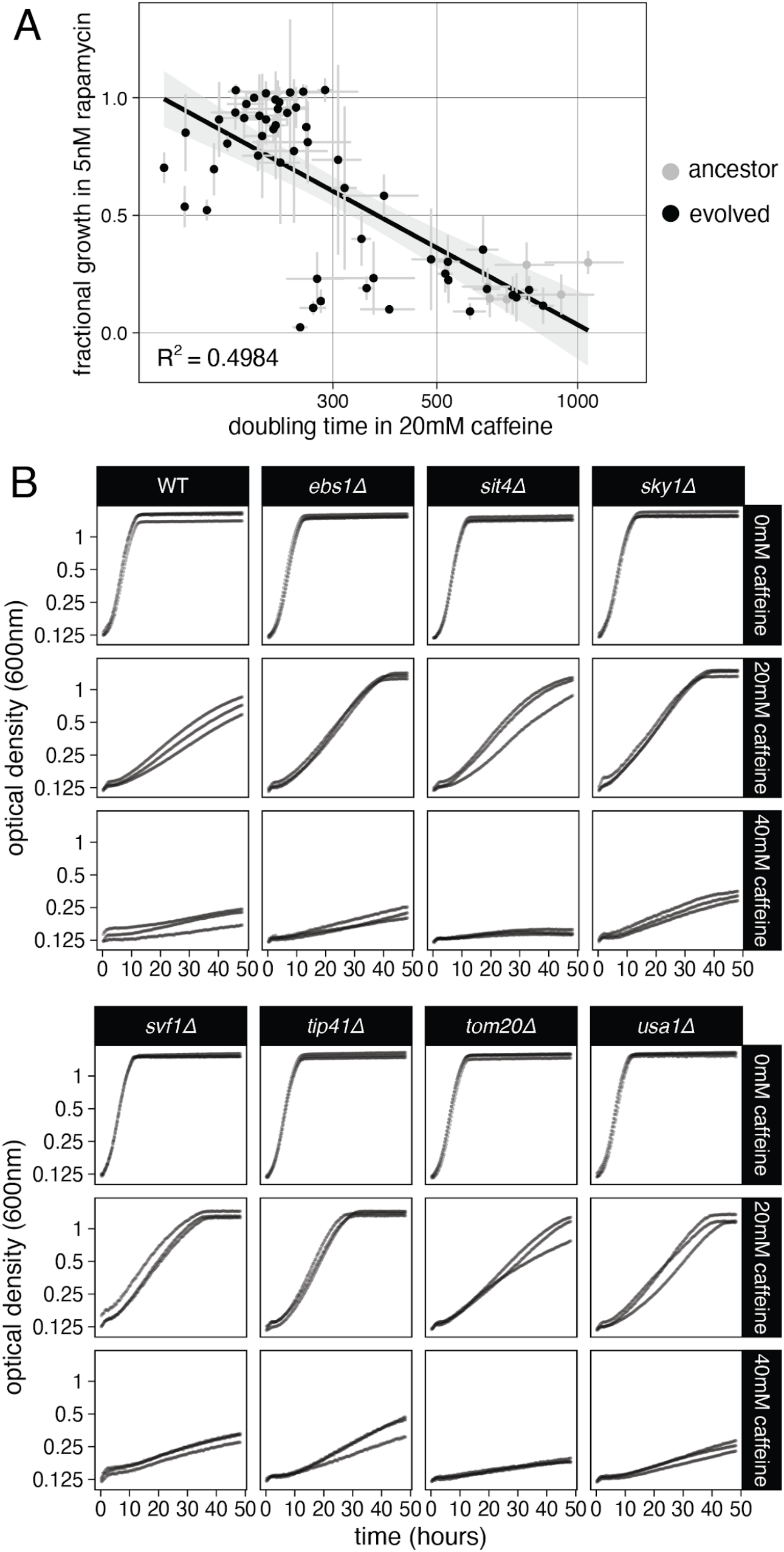
(A) Relative growth of clones in presence vs. absence of 5nM rapamycin, measured by optical density after 24 hours compared to growth in caffeine. (B) Growth of strains with indicated deletions in caffeine media.

**Supplemental Figure 5:**
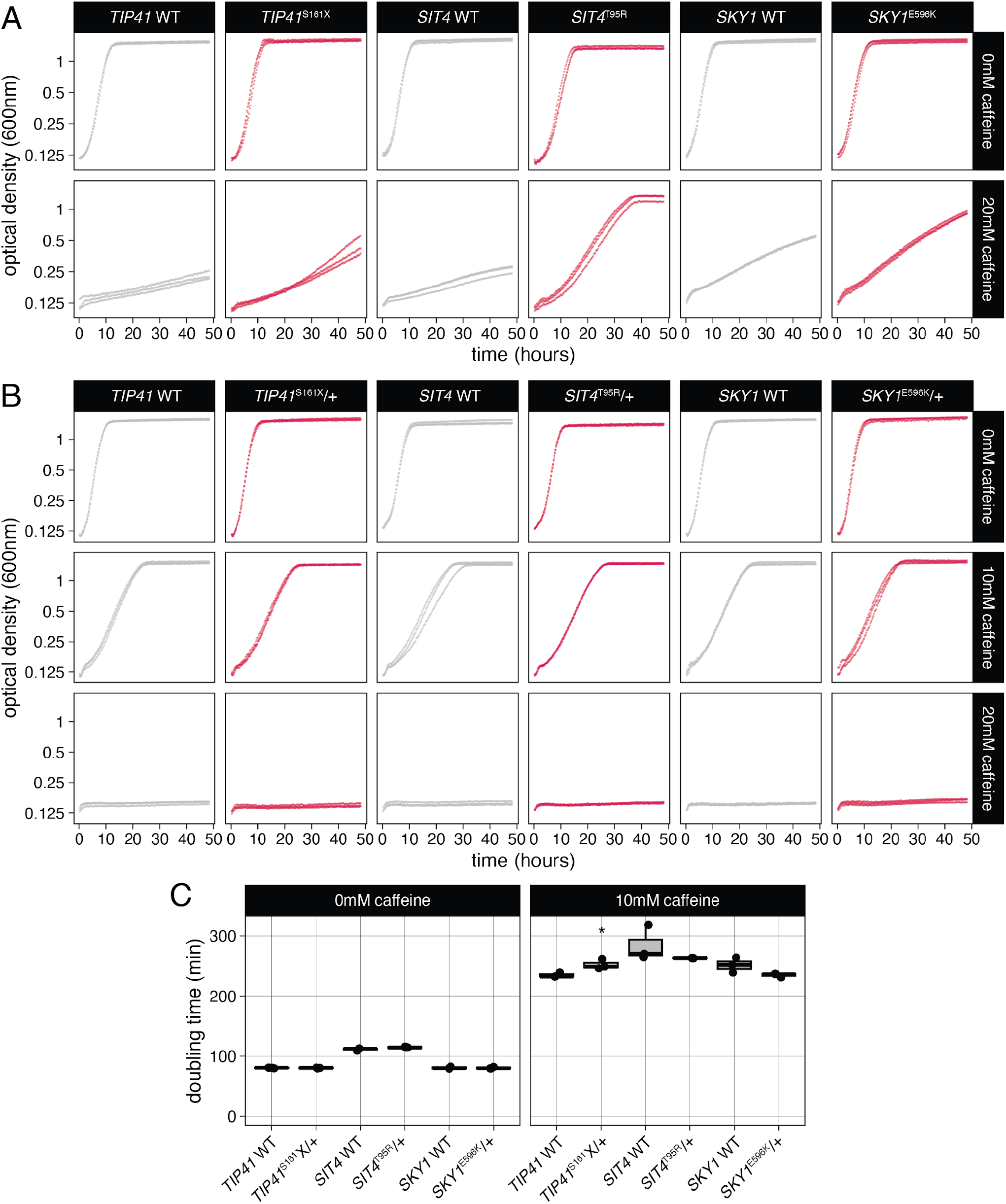
(A) Growth of haploid and (B) heterozygous diploid strains with indicated mutations in media containing caffeine. Each contains a synonymous mutation, see Table S1 for details. (C) Doubling time of diploid strains from B.

